# From Biofilms to Biocontrol: Aquaculture Biofilms as a Reservoir of Pseudomonas spp. with Activity Against Fish and Human Pathogens

**DOI:** 10.1101/2025.11.05.686791

**Authors:** Jennifer N.T. Nguyen, Todd Testerman, Kathryn R. McBride, Hailey Donohue, Jeremiah Marden, Marcy J. Balunas, Joerg Graf

## Abstract

The antimicrobial resistance crisis, driven by excessive use of antimicrobials in medical and agricultural settings, has spurred the search for alternative strategies. In the present study, we investigated surface biofilms from a rainbow trout aquaculture facility to identify bacteria with inhibitory activity against fish and human pathogens. A total of 906 isolates were obtained, of which 478 were taxonomically classified using full-length 16S rRNA gene sequencing, revealing *Pseudomonas*, *Aeromonas*, and *Flavobacterium* as the most abundant genera. Twenty-nine isolates, predominantly novel *Pseudomonas* species, inhibited one or more fish pathogen strains, with two *P. aphyarum* strains inhibiting every fish pathogen tested. All human pathogen strains were inhibited by at least one trout farm isolate. A biofilm assay identified strains capable of excluding *F. columnare* from established biofilms. Numerous strains reduced biofilm colonization, with *P. aphyarum* and *P. idahonensis* strains exhibiting the greatest reduction. Genome analysis and biosynthetic gene cluster (BGC) identification from a subset of isolates revealed shared and unique clusters with strong potential for antimicrobial production, correlating in part with observed inhibition patterns. Metabolomics profiling identified a suite of strain-specific siderophores that may mediate biofilm disruption. Additionally, co-culture of these isolates with the human pathogen *Pseudomonas aeruginosa* enhanced the production of specialized metabolites with reported antimicrobial activities. Collectively, these findings highlight trout farm biofilms as a promising source of bacteria that could be developed as probiotics or investigated for novel antimicrobial discovery.

**Importance:** Disease outbreaks in aquaculture facilities pose significant threats to food security and often rely on antibiotic treatments, contributing to antimicrobial resistance. This study identifies *Pseudomonas* isolates from native biofilms of a trout aquaculture facility that are capable of disrupting biofilms and inhibiting a wide range of fish and human pathogens. These *Pseudomonas* isolates produce a variety of siderophores that may mediate biofilm disruption, offering a promising avenue for developing probiotic applications or the discovery of new antimicrobials to improve aquaculture health and reduce reliance on antibiotics. The discovery of these biofilm-disrupting bacteria holds potential significance for combating antimicrobial resistance in both aquaculture and clinical settings.

## Introduction

Members of microbial biofilms proliferate in a hotbed of competition, with species constantly competing for space and nutrients (1, 2). The competition between microbes can be divided between exploitive competition (indirect interactions such as competition for resources) and interference competition (direct or antagonistic interactions) (3). Numerous interference mechanisms are used by these microbes and have been extensively studied including siderophores (4), type VI secretion systems (5), bacteriocins (5), and quorum sensing inhibition (6), among others (7, 8). The organisms existing within a biofilm structure are in close proximity and enclosed by matrix composed primarily of exopolysaccharides (9). These biofilm communities are incredibly resilient to chemical and physical treatments and facilitate interactions between microbial taxa, including horizontal gene transfer events, promoting the development of antibiotic-resistant organisms (10–12).

In freshwater systems such as rivers and aquaculture settings, the surface biofilm communities are complex, consisting of bacteria, fungi and microalgae (13, 14). In temperate environments, bacterial communities are often dominated by the classes Alphaproteobacteria, Bacteroidia, Cyanobacteria, Gammaproteobacteria, Planctomycetes, and Verrucomicrobia (14, 15). These microbial communities perform important functions including nutrient recycling and biodegradation of various compounds as well as microplastics (16), but they can also harbor human or fish pathogens (15, 17, 18).

Aquaculture, the culture and harvest of aquatic species such as shrimp, shellfish and fish, has been a consistently growing food production sector for the last 30 years (19). However, a major challenge to this continued growth has been bacterial diseases which can significantly reduce total production capacity (20, 21). With respect to the largest worldwide sector of aquaculture, finfish production (22), production capacity is substantially limited by bacterial pathogens such as *Aeromonas salmonicida*, *Yersinia ruckeri*, *Flavobacterium columnare* and *F. psychrophilum* (23). Preventative and curative measures such as vaccination and antibiotics can be effective but frequently fail to stem the problem. Vaccination can have limited effectiveness depending on the antigenic variability of the pathogen (24, 25), host immune system structure (26), and logistical considerations for vaccinating the farmed species (27). Antibiotics can be used in response to an outbreak, but their overuse in a prophylactic capacity has led to increased resistance to many antibiotics for several bacterial pathogens (28, 29).

In the present study, we sought to source and characterize bacteria isolated from surface biofilms at a freshwater rainbow trout (*Oncorhynchus mykiss*) aquaculture facility. Postulating that biofilm bacteria likely produce numerous secondary metabolites and antimicrobials, we designed a strategy to identify constituents of the native biofilm communities that may prevent pathogenic bacteria from colonizing. The bacteria isolated from these surface biofilms were identified using full-length 16S rRNA PacBio gene sequencing and then screened for antagonistic activity against economically impactful fish pathogens, as well as common human pathogens. Genome mining techniques were then employed to identify biosynthetic gene clusters (BGCs), and untargeted metabolomics was used to profile metabolites produced by selected strains. Lastly, these strains were co-cultured with a human pathogen to identify small molecules that may contribute to the observed antibiotic and biofilm inhibition properties. Overall, we were interested in determining if strains from an aquaculture facility would be able to inhibit the growth of pathogens and be a potential tool for excluding pathogenic bacteria in biofilms.

## Materials and Methods

### Sampling Site

The isolates were collected from a flow-through, spring-fed, freshwater rainbow trout aquaculture facility in Idaho, USA. The primary sampling site was the indoor hatchery facility, and a secondary sampling site was the natural spring that provides the water for the farm located within Box Canyon State Park. Within the hatchery, 20 separate epoxy-coated concrete raceways contain the maturing fish, with water from the Box Canyon spring (BCS) entering and then being discharged into the nearby river. The BCS site contains wild populations of freshwater fish and receives abundant sunlight. The isolates were collected in October of 2019. The water temperature at the time of sampling at both sites was 15°C. The outflowing water from the hatchery had the following water quality metrics: 0.074 mg/L total phosphorus, 7.8 mg/L dissolved oxygen, < 31 mg/L chloride, 0.5 mg/L ammonia and pH 7.3.

### Isolate Collection

Approximately 1-meter-long strips from the surfaces of the walls and rocks at the hatchery and BCS, respectively, were swabbed using sterile Whatman OmniSwabs (Maidstone, United Kingdom). Individual fish were caught in a net, and a sterile swab was run over the exterior of each animal (this was non-lethal). These swabs were then streaked onto Petri plates of microbiological media. To account for potential overcrowding on culture plates hindering our ability to obtain pure cultures, a new swab was then inoculated on the initial plate and then streaked onto a second Petri plate to dilute the overall culture. This process was repeated with a new swab and a new plate resulting in one “primary” plate and two “dilution” plates. The media types used were Reasoner’s 2A agar (R2A) (DSMZ #830), tryptic soy agar (TSA) (BD, Franklin Lakes, NJ, USA), 1:100 diluted TSA (BD), nutrient agar (BD), or Aeromonas agar (Millipore- Sigma, Burlington, MA, USA). Inoculated plates were stored at 4°C until being shipped overnight in an insulated cooler with ice packs.

### Isolation

Upon receipt, plates were incubated at either 15 or 25°C. Isolated colonies were harvested from these plates over a period of 2 months (to account for fast and slow growing bacteria) and subsequently streaked out onto a fresh plate to ensure purity of the culture. A single colony from these subcultured plates was then subcultured again onto a fresh plate. Purity was assessed visually, and isolates deemed pure were then scraped off the plate, resuspended in their corresponding broth medium with a final concentration of 20% glycerol, and frozen at -80°C. If an isolate still did not achieve visual purity, additional subcultures were performed until purity was established.

### Identification

Isolates were identified by sequencing the nearly full-length 16S rRNA gene using PacBio HiFi sequencing technology. For the identification of isolates, strains were chosen from the early, middle, and late growers (the first, second and third 33% isolated, respectively) and inoculated from frozen stock directly into six 96-well, deep-well plates containing 600 µL liquid R2 medium (Greiner Bio-One, Monroe, NC, USA; cat # 780261). These plates were incubated for 48 h at 19°C with shaking at 200 rpm, after which 10 µL of dense broth culture was used for DNA extraction and purification with the Shoreline Biome V1-V9 sequencing kit as previously described (Shoreline Biome, Farmington, CT, USA) (30). PCR amplification of the full-length 16S rRNA gene was performed using the Shoreline Biome V1-V9 sequencing kit with the following thermocycler conditions: 95°C for 60 s followed by 35 cycles of 95°C for 30s, 63°C for 45 s (ramp speed of 4°C/s), and 72°C for 90 s, with a final extension step of 72°C for 180 s. A QIAxcel Advanced System (Qiagen, Hilden, Germany) was used to confirm reaction success as indicated by a band near 1,500 bp. PCR reactions were then pooled using equal volumes from each reaction and cleaned using the Qiagen GeneRead Size Selection Kit (Qiagen). Cleanup success was verified by running the cleaned, pooled libraries on an 0.8% agarose gel and verifying the absence of primer dimer and other extraneous bands. From each 96-well plate, one SMRT-bell library with a different bar code was prepared. Subsequently, the libraries were combined and sequenced on a PacBio Sequel (Pacific Biosciences, Menlo Park, CA, USA) and processed for circular consensus sequencing (CCS) reads.

The sequencing reads were processed using the SBAnalyzer software (version 3.0) along with DADA2 (version 1.16) (31) to call amplified sequence variants, ASVs. Briefly, CCS reads were demultiplexed into separate FASTQ files using SBanalyzer. Demultiplexed CCS reads were trimmed and oriented using dada2::removePrimers (31). Trimmed CCS reads were filtered using dada2::fastqFilter (31). Filtered CCS reads were separated manually into bins based on their read length in R (version 3.6) (32). Binned CCS reads were used as input for ASV inference using dada2::dada (31). Per-bin ASV inference results were merged in R to produce per-plate ASV inference results. Merged per-plate results were merged in R to produce the final outputs (frequency table and ASV FASTA file). Following ASV generation, abundance tables and representative sequences were imported into QIIME2 (version 2020.2) (33). The data was filtered with a requirement that an isolate needed a minimum of 200 reads for inclusion, and each individual ASV needed 200 reads summed across all isolates for inclusion. Taxonomy assignments were performed in QIIME2 using the VSEARCH consensus plugin with the All-Species Living Tree Project (LTP) reference database (version 132) (34). The final taxonomic assignment per isolate used the highest percent identity value hit with a minimum of 97% identity required for species level assignment. Trees were generated by extracting all reads for the genus of interest from the LTP reference database and the isolate dataset. Reads were then combined, and a tree was constructed using the align-to-tree-mafft-fasttree plugin in QIIME2.

The resulting rooted tree was then imported into iTOL (version 6.5.8) (35) for annotation and figure generation.

### Genome analysis

The genomes of six *Pseudomonas isolates* (*P. fontis* isolate 656, *P. rubra* isolate 291, *P. aphyarum* isolates 233, 386, and 387, and *P. idahonensis* isolate 357) were retrieved from NCBI (BioProject <u>PRJNA835116</u>). Genomes were annotated with Prokka v1.14.6 (36), then functionally categorized with eggnog-mapper v2 (37). Pangenomic analysis was performed with panaroo v.1.2.7 (38) using default parameters. BGC prediction was performed with antiSMASH v7 (39) using default relaxed setting parameters. Clustering of BGCs was performed with BiG-SCAPE v1.1.5 (40) using the default settings, specifying a distance cutoff of 0.5 for clustering with auto clustering mode.

### Plate Inhibition Assays

Fish and human pathogens were inoculated into R2 broth (fish pathogens) or nutrient broth (human pathogens). All pathogen strain information is available in Table S1. Following 24 h of growth, a cotton swab was soaked in the resulting culture and used to spread a lawn of pathogen on a single-well agar plate (Thermo Scientific, Waltham, MA, USA; cat # 242811) of the same media type. Inoculated deep-well plates containing the biofilm isolates (as described in the “Identification” section of the Materials and Methods) were incubated for 48 h at 19°C with shaking at 200 rpm. Following incubation, the pathogen plates and an isolate plate were loaded into an epMotion 5073 liquid handling robot (Eppendorf, Hamburg, Germany). Following mixing, 2 µL of each biofilm isolate was spotted onto the pathogen lawn, with 48 isolates spotted onto each plate. These plates were then incubated until a visible lawn was present (24-48 h) at 19°C (fish pathogens) or 25°C (human pathogens) and visually observed for zones of inhibition.

### Biofilm Inhibition Assays

The 29 bioactive isolates from the plate inhibition assays were cultured overnight in Shieh medium (41) at 30°C. A *Flavobacterium columnare* strain isolated from a previous columnaris outbreak at this specific aquaculture facility was introduced with a GFP-expressing plasmid harboring a tetracycline resistance marker (pNT67) via conjugation with S17-1 λpir (42). This strain was not used in the initial plate inhibition assays as it did not grow in R2A medium. For the biofilm-inhibition assay, this strain was streaked out from frozen stock onto Shieh agar supplemented with 5 µg/mL tetracycline and incubated for 24 h at 30°C. Multiple colonies were then picked and inoculated into liquid Shieh medium (without antibiotic) in a baffled flask (Fisher, cat # 09-552-34) and incubated at 30°C for 24 h with shaking at 200 rpm. Isolate cultures were diluted in sterile Shieh broth to approximately 0.08-0.15 OD600, after which 500 µL of each dilution was transferred into a 48-well sterile culture plate (Corning, Corning, NY, USA, cat # 351178). Culture plates were then incubated statically at 30°C for 48 h. Following incubation, the supernatant was removed, the well was washed once with 500 µL of sterile Shieh broth, and then 500 µL of the *F. columnare* GFP strain was inoculated into each well at approximately 0.08-0.15 OD600. The culture plate was then measured on a plate reader (SpectraMax i3x, Molecular Devices, San Jose, CA) to measure the initial OD600 and GFP fluorescence and then incubated statically at 30°C for 72 h to allow sufficient time for *F. columnare* growth. Following incubation, the culture plate was again read on a plate reader to measure the final OD600 and GFP fluorescence. The supernatant from each well was then removed, the well washed once with 500 µL of sterile Shieh broth, and then 500 µL of phosphate buffered saline (PBS) was added to resuspend the biofilm. The plate was then incubated for 1 h at 30°C with shaking at 200 rpm to allow time for resuspension. The plate was then read on the plate reader to measure the final OD600 and GFP fluorescence of the biofilm fraction. Kruskal-Wallis testing was used to compare biofilm inhibition results and Dunn’s multiple comparison test was used for *post-hoc* pairwise comparisons.

### Large Scale Cultures and Co-cultures with Pseudomonas aeruginosa

Cultures and co-cultures were prepared using previously described methods (43) with the following modifications. *Pseudomonas* isolates from the trout facility biofilms (strains 233, 386, 387, 291, 357, and 656) were grown on YSP agar for 24 h at 30°C. On day zero, liquid seed cultures of each *Pseudomonas* isolate were prepared by adding one colony into 5 mL of NB liquid media in 6 deep-wells plate, which were then incubated for 48 h at 30°C with shaking at 200 rpm. On the same day, the human pathogen *Pseudomonas aeruginosa* (ATCC 15442) was streaked onto YSP agar for 24 h at 30°C. On day two, 1.5 mL of each *Pseudomonas* isolate seed culture was added to 50 mL of NB liquid media in separate 125-mL flasks, one for monoculture and one for co-culture, and incubated for 24 h at 30°C with shaking at 200 rpm. Concurrently on day two, one colony of *P. aeruginosa* was inoculated into 5 mL of NB liquid media in a 6-well deep-well plate and incubated for 24 h at 30°C with shaking at 200 rpm. On day three, 400 μL of *P. aeruginosa* was added to the respective co-culture flask, and all flasks were incubated for an additional 24 h at 30°C with shaking at 200 rpm. On day four, cultures were harvested for use in metabolomics.

### Metabolomics

#### Sample extraction

All solvents were of high-performance liquid chromatography (HPLC) grade and purchased from Sigma Aldrich. Liquid cultures of bacterial strains in 50 mL of NB media were sonicated for 60 s and extracted with an equal volume of ethyl acetate three times. The ethyl acetate layer was collected after each extraction, combined, dried using rotary evaporation, and extracts were transferred to vials and stored at -80°C.

#### Mass spectrometry (MS) data acquisition and processing

Extracts were prepared at 1 mg/mL in 50% methanol (MeOH). Data were acquired using a Bruker timsTOF Pro2 (Bruker-Daltonics, Billerica, MA, USA) coupled to an Agilent 1290 Infinity II Bio UHPLC (Agilent, Santa Clara, CA, USA) using an Acquity UPLC HSST3 column (2.1 × 150 mm, 1.8 μM). Each sample (2 μL) was injected in technical triplicate at random. Samples were eluted using a 0.4 mL/min gradient of mobile phases A (0.1% formic acid in water) and B (0.1% formic acid in acetonitrile) employing the following conditions: 1 min hold at 5% B, 1 min ramp to 15% B, 6 min ramp to 100% B, 3 min hold at 100% B, 0.1 min ramp back to 5% B and a re-equilibration hold at 5% B for 1.4 min.

Data acquisition was performed in positive ionization mode using an ESI source, with a collision energy of 10 eV, capillary voltage of 4500 V, dry temperature of 220°C, sheath gas temperature of 220°C, mass range of 50-2000 *m/z*, and mobility (1/Ko) range of 0.45-1.45 V⋅s/cm^2^. Fragmentation data were acquired with a collision energy of 50 eV, with 2 PASEF MS/MS scans per cycle, for a total cycle of 0.53 s.

Once acquired, MS data were preprocessed using Bruker MetaboScape^®^ version 9.0.1 (Bruker-Daltonics, Billerica, MA, USA) using the MCube T-Rex 4D Metabolomics workflow for peak picking and alignment. The intensity threshold was experimentally determined by comparison of the baseline noise from samples and 50% MeOH blanks, resulting in a baseline intensity threshold of 500 counts. The resulting feature table was then processed using mpactR (44) with the following parameters: mispicked peak correction - ringing mass window of 0.5 atomic mass units (AMUs), isotopic mass window of 0.01 AMU with a maximum isotopic mass shift of 3 AMUs, and a tR window of 0.05; in-source ion filtering threshold of 0.95 Spearman correlation; median coefficient of variation (CV) of technical replicates of 0.5; and blank filtering using 50% MeOH blanks at a 0.05 threshold.

For *in-silico* formula prediction and annotations, MetaboScape^®^, NPAtlas (45), and SIDERITE (46) were used as databases for annotations, with a 5 ppm upper limit for formula prediction and a 10 ppm upper limit for annotation. Annotations were further verified by comparing fragmentation patterns using public data [GNPS: Global Natural Products Social Molecular Networking database (47), MoNA, and MassBank (48)] when available, or using the Competitive Fragmentation Modeling for Metabolite Identification (CFM-ID) spectra prediction (49).

#### Multivariate analysis

The R package mpactR (44) was used to conduct statistical comparisons between metabolomes of monocultures and co-cultures, visualized as heatmaps and volcano plots. The heatmap was built using normalized raw counts within each feature between minimum and maximum counts for that feature. Samples were grouped on the x-axis by overall metabolomic similarity, with features grouped on the y-axis. The volcano plot was generated using multiple two tailed t-tests, including false discovery rate (FDR) correction using the Benjamini-Hochberg procedure (50).

### 96-well Antibacterial Bioassays

Extracts from monocultures and co-cultures of *Pseudomonas* isolates were tested for antibacterial activity against the human pathogen methicillin-sensitive *Staphylococcus aureus* (MSSA; strain TCH 959). Bioassays were performed in 96-well flat bottom plates (Corning Costar TM, Kennebunk, ME, USA) using Tryptic Soy Broth (BD Difco™, Sparks, MD, USA), incubated at 30°C.

Bioassays were performed as previously described (51) in 96-well plates (Corning Costar, Corning, NY, USA) with the following modifications: MSSA pathogen inoculum was adjusted to OD600 0.1 [(approximately 1-2 × 10^8^ CFU/mL (52)] prior to use. A master mix was prepared using 1.6 mL diluted MSSA bacterial inoculum, 7.84 mL sterile water, and 6.4 mL of broth. Master mix was aliquoted into each well (199 μL) to which was added 1 μL of either positive control (vancomycin, final testing concentration of 2.5 μg/mL), negative control [dimethyl sulfoxide (DMSO)], or extract prepared in DMSO (screened at final concentration of 500 μg/mL). Sterility was measured using wells containing 99 μL of sterile water, 100 μL of broth, and 1 μL of DMSO. Samples and controls were tested in triplicates. Plates were read at 0 and 24 h at 600 nm using a Synergy HT (Biotek, Winooski, VT, USA). Results were normalized to the DMSO negative control to calculate percent control activity (PCA).

## Results

### Isolation and Identification of Biofilm Isolates

Surface biofilms were swabbed from 6 hatchery raceways, 4 locations within BCS, and 3 separate juvenile fish within the hatchery. A total of 906 isolates were stocked, of which 510 isolates were processed for identification (Fig. 1, Table S2). Of these 510 isolates, 478 isolates returned a predominant taxon (≥60% of reads belonging to a single genus) based on the 16S rRNA gene sequencing results. The per-genus isolate counts are presented in Table S3. Species assignment of these isolates to their most closely related type-strain sequence indicated that a minimum of 122 species were recovered (Table S3). The major genera comprising 75% of all identified isolates were *Pseudomonas* (28%), *Aeromonas* (18%), *Flavobacterium* (13%), *Acinetobacter* (7%), *Brevundimonas* (4%), *Microbacterium* (3%), and *Sphingopyxis* (2%) (Fig. 2). In total, 55 genera were identified spanning 5 bacterial phyla.

**Figure 1.**
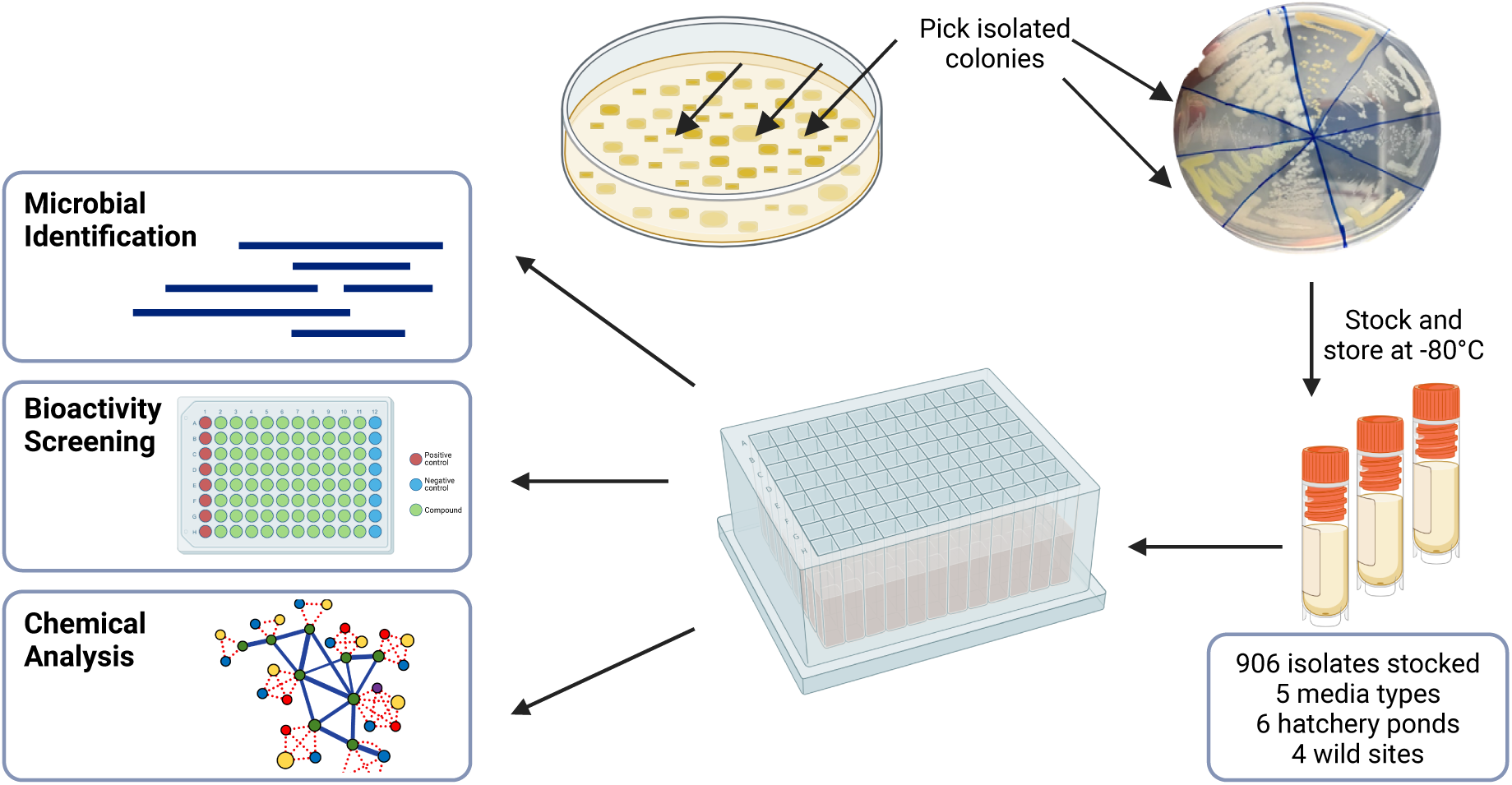
Isolate collection, identification, screening, and chemical analysis. In total, 906 isolates were isolated from the aquaculture facility after incubation. Of these isolates, 510 were identified via 16S rRNA sequencing and screened against a panel of fish and human pathogens. Six bioactive isolates were selected for chemical analysis. Figure created using BioRender.

**Figure 2.**
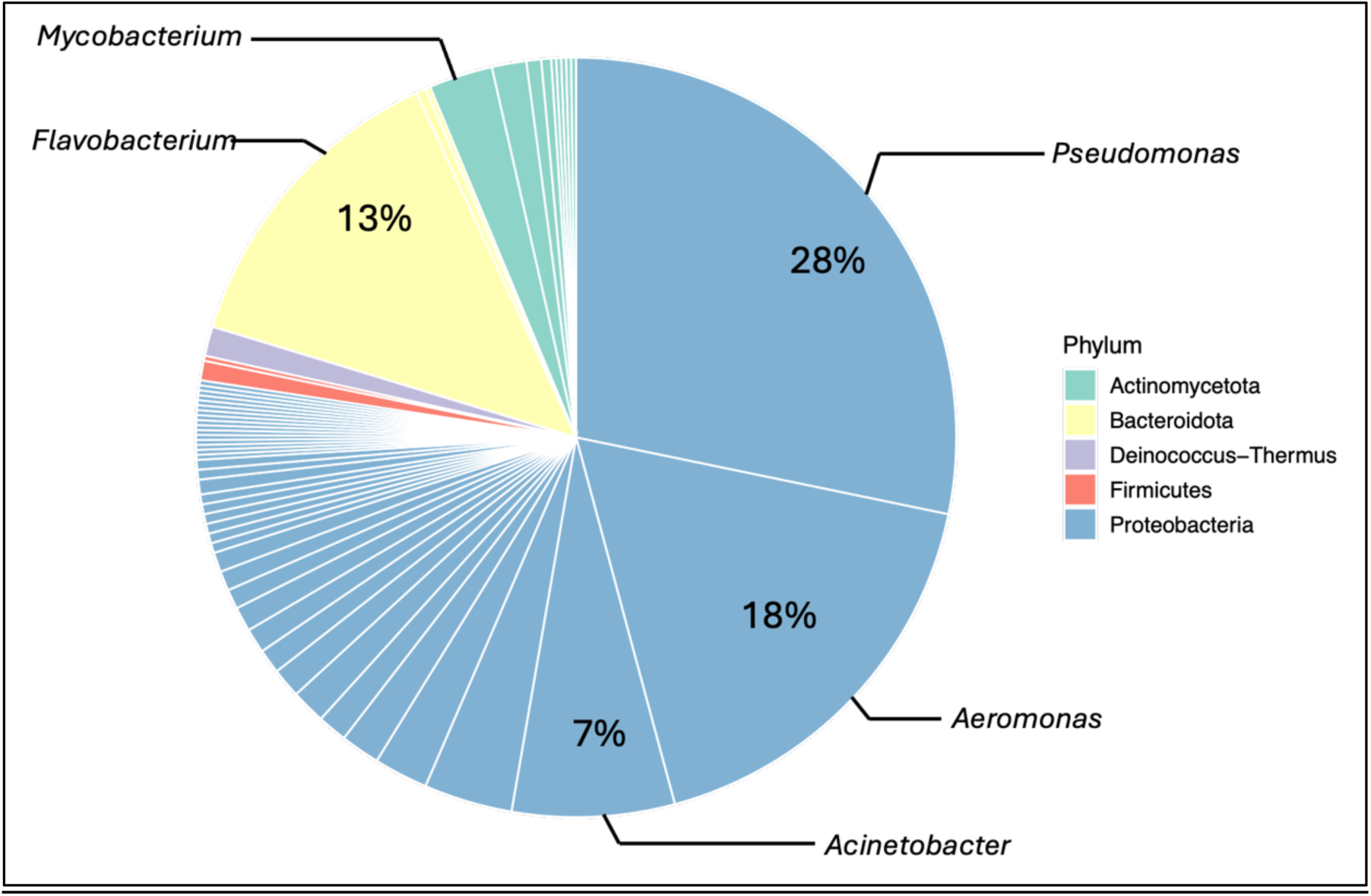
Taxonomic summary of identified isolates. Surface biofilm bacterial community members were identified using full-length 16S rRNA PacBio gene sequencing. In total, 55 genera were identified spanning 5 bacterial phyla with a minimum of 122 species based on most closely related type strain sequence. The sampling revealed that Proteobacteria was the most prominent taxon cultured from the aquaculture biofilm community, with 28% of isolates belonging to the *Pseudomonas* genus. Comprehensive taxonomic profiling of the identified isolates is summarized in Table S3.

To assess relative species diversity of the recovered isolates within these genera, 16S rRNA gene phylogenies were produced from all ASVs and available reference type strain sequences for the genus of interest (Fig. S1-7). These phylograms show that a wide diversity of species groups were recovered, with large clades being formed by some isolate groups with high bootstrap support, possibly indicating numerous new species.

### Inhibition of Pathogens by Aquaculture Biofilm Isolates

Initial pathogen inhibition screening included the type strains of four economically relevant fish pathogens: *F. columnare*, *F. psychrophilum*, *A. salmonicida*, and *Y. ruckeri*. Isolates identified as inhibitory during this initial screening were subsequently screened against an expanded panel including multiple strains and subspecies from these four species along with other fish pathogens. Of the 29 isolates identified as inhibitory during the initial screens, 26 were *Pseudomonas* spp., 2 were an *Aeromonas* sp., 1 was a *Serratia* sp., 1 was a *Microbacterium* sp., and 1 was a *Janthinobacterium* sp. (Fig. 3A, Table S8). Subsequent genome sequencing of some isolates allowed species-level grouping and assignment (Table S4), while the remaining isolates were grouped into species groups solely based on the 16S rRNA gene data.

**Figure 3.**
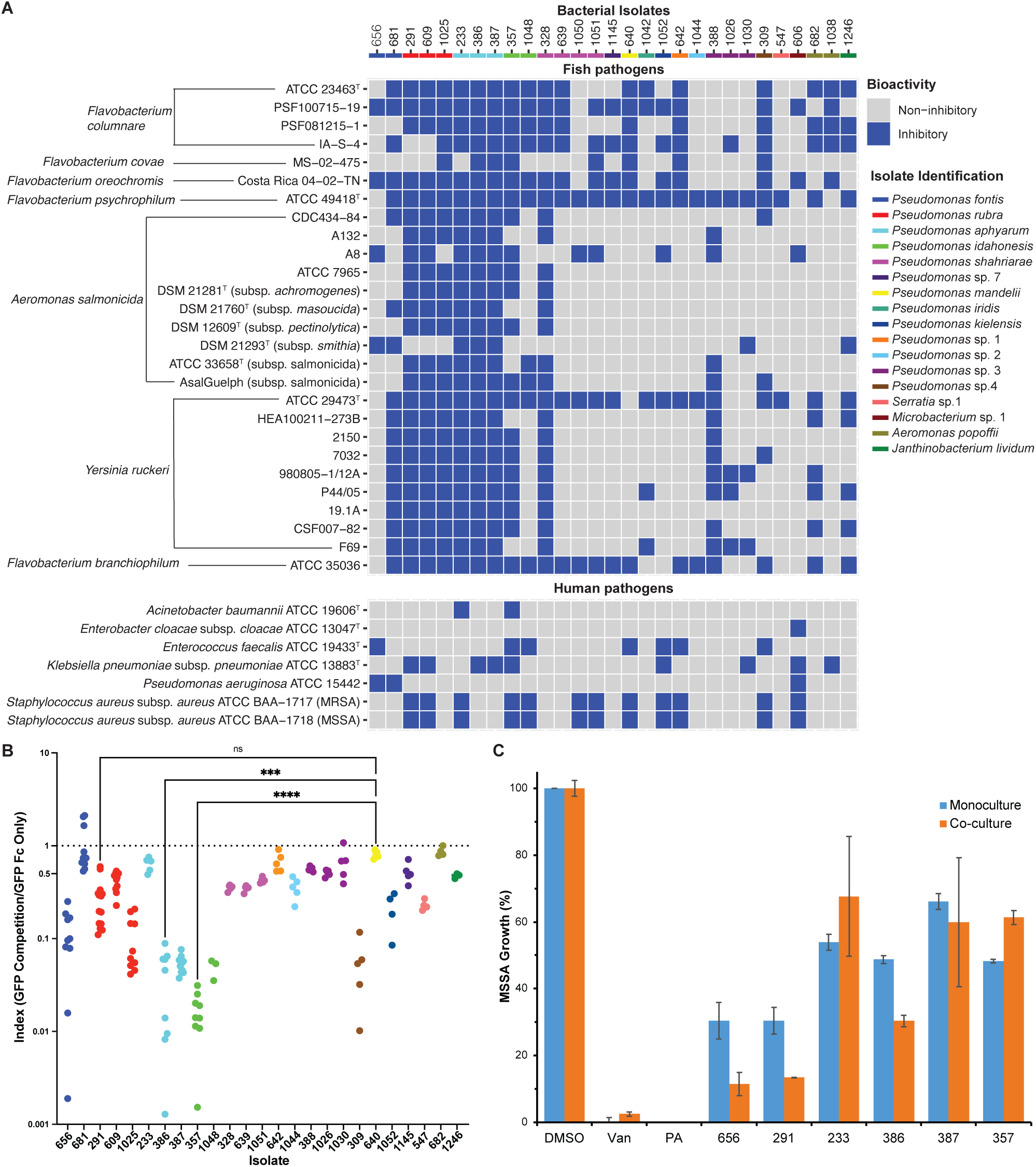
Surface biofilm bacterial isolates exhibit inhibitory activity against fish and human pathogens and can exclude a relevant fish pathogen from forming biofilms. (**A**) In total, 29 surface biofilm bacterial isolates exhibited inhibitory activity against a panel of fish and human pathogens in plate inhibition assays. The majority of the inhibitory isolates belong to the genus *Pseudomonas.* (**B**) A subset of those isolates excluded *F. columnare* from colonizing biofilms in a biofilm exclusion assay. (**C**) Co-culture of each of the six bioactive isolates from the biofilm inhibition assay with *P. aeruginosa* yielded extracts with increased antibacterial activity against methicillin-sensitive *Staphylococcus aureus* (MSSA) for a subset of isolates, including strains 656, 291, and 386 [Van = vancomycin (positive control), PA = *P. aeruginosa* extract, and DMSO (negative control)].

In our initial screens, many aquaculture isolates exhibited broad activity against fish pathogens and human ESKAPE pathogens. *Pseudomonas rubra* (isolates 291, 609, and 1025), *Pseudomonas aphyarum* (233, 386, and 387) and one isolate of *Pseudomonas shahriarae* inhibited between 24/27 and 27/27 fish pathogens (*A. salmonicida*, *Flavobacterium* spp., and *Y. ruckeri*) tested (Fig. 3A). The remaining *Pseudomonas* species, as well as the non-*Pseudomonas* inhibitors [*Serratia* sp. 1 (547), *Microbacterium* sp. 1 (606), *Aeromonas popoffii* (682 and 1038), and *Janthinobacterium lividum* (1246)], were much less effective at inhibiting the fish pathogens used in our study. Interestingly, *F. covae* was seemingly resistant to inhibition compared to the other *Flavobacterium* strains, including the *F. columnare* strains that until recently included *Flavobacterium covae* and *Flavobacterium oreochromis* (53). *F. psychrophilum* was inhibited by 26/29 biofilm strains. Overall, we succeeded in isolating seven strains that were highly active across all fish pathogens tested.

A second series of screens was subsequently undertaken using the hospital-associated ESKAPE pathogens (*Enterococcus faecium*, *Staphylococcus aureus*, *Klebsiella pneumoniae*, *Acinetobacter baumannii*, *Pseudomonas aeruginosa*, and *Enterobacter* spp.) relevant in human healthcare settings (Fig. 3A, Table S9). The isolates inhibiting the greatest number of pathogenic strains were isolates 357 (*Pseudomonas idahonensis*) and 606 (*Microbacterium* sp. 1), each inhibiting 5/7 tested strains. Additionally, both isolates from species group *P. fontis* (656 and 681) inhibited *P. aeruginosa*, a well-studied gram-negative human pathogen with biofilm formation as a known virulence mechanism (54). An additional 6/29 tested isolates inhibited all the gram-positive pathogens tested, namely *E. faecalis* and the two *S. aureus* strains.

The twenty-five *Pseudomonas* isolates used in the plate inhibition assay were subsequently tested for their ability to exclude a pathogenic *F. columnare* strain, CSF-298-10, from a preformed biofilm. CSF-298-10 was originally isolated from the same farm that the inhibitory isolates were sourced from (55) and was introduced with a GFP-expressing plasmid. Five isolates strongly inhibited the ability of *F. columnare* to colonize the biofilms, including 357 (*P. idahonensis*), 386 (*P. aphyarum*), 387 (*P. aphyarum*), 1048 (*P. idahonensis*), and 309 (*Pseudomonas* sp. 4) (Fig. 3B, Table S10). Within species-group variation was interesting to note, with isolate 681 performing far worse than isolate 656 (*P. fontis* group), and isolate 233 performing far worse than isolates 386 and 387 (*P. aphyarum* group) (Fig. 3B).

### Bacterial metabolites from aquaculture strains exhibit antibacterial activity

To identify bacterial metabolites that might be responsible for the antimicrobial and/or biofilm inhibitory activity, a subset of six isolates (strains 656, 291, 233, 386, 387, and 357) were selected for further investigation. *P. fontis* 656 was selected based on activity against *P. aeruginosa*. *P. rubra* 291 was selected based on broad antimicrobial activity against the fish pathogens. *P. aphyarum* 233, 386, and 387 were selected based on intra-species differential activity against the human pathogens and in the biofilm exclusion assay. *P. idahonensis* 357 was selected for its significant activity in the biofilm exclusion assay.

Extracts of these six isolates were tested against MSSA, with isolates 656 and 291 exhibiting the strongest antibacterial activity (70% inhibition), while isolates 386 and 357 exhibited moderate antibacterial activity (50% inhibition) (Fig. 3C). Differential inhibitory activity was not observed between extracts of *P. aphyarum* isolates (233, 386, and 387), as had been shown in the initial screen. Because these bacterial isolates originate from a biofilm system where there is significant contact and cross-species communication, we also co-cultured each individual isolate with a small inoculum of *P. aeruginosa* to elicit secondary metabolism. Co-culture of each isolate with *P. aeruginosa* increased the antibacterial activity in 3/6 isolates (Fig. 3C), including strains 656, 291, and 386, which showed significantly stronger activity against MSSA (an additional 19, 17, and 18% inhibition, respectively).

### In Silico Investigation of Biosynthetic Potential

To investigate possible mechanisms behind their differential inhibitory phenotypes, an *in silico* approach was used with the subset of six bioactive *Pseudomonas* isolates for which we had previously sequenced genomes (56). Phylogenetic relationships of proteins encoded in the genomes of these six isolates were analyzed using the Cluster of Orthologous Genes (COGs) database (57), enabling functional assignments for proteins that may have not been studied experimentally, which is particularly relevant when investigating environmental isolates.

Comparative pangenome analysis with panaroo (38) identified 9670 orthologous genes across these six genomes. Notably, 74% of these genes were present in less than half of the isolates (< 3 isolates, represented in orange; Fig. 4A), indicating considerable genomic diversity. Genes associated with secondary metabolite biosynthesis and transport were detected across all isolates, consistent with their observed antimicrobial activity (Fig. 3) and supporting their potential for bioactive metabolite production. However, the majority of secondary metabolite proteins were shared between three or fewer isolates, suggesting that these bacterial isolates hold diverse metabolic roles in the biofilm community and may produce diverse chemistry to form and sustain the biofilm environment.

**Figure 4.**
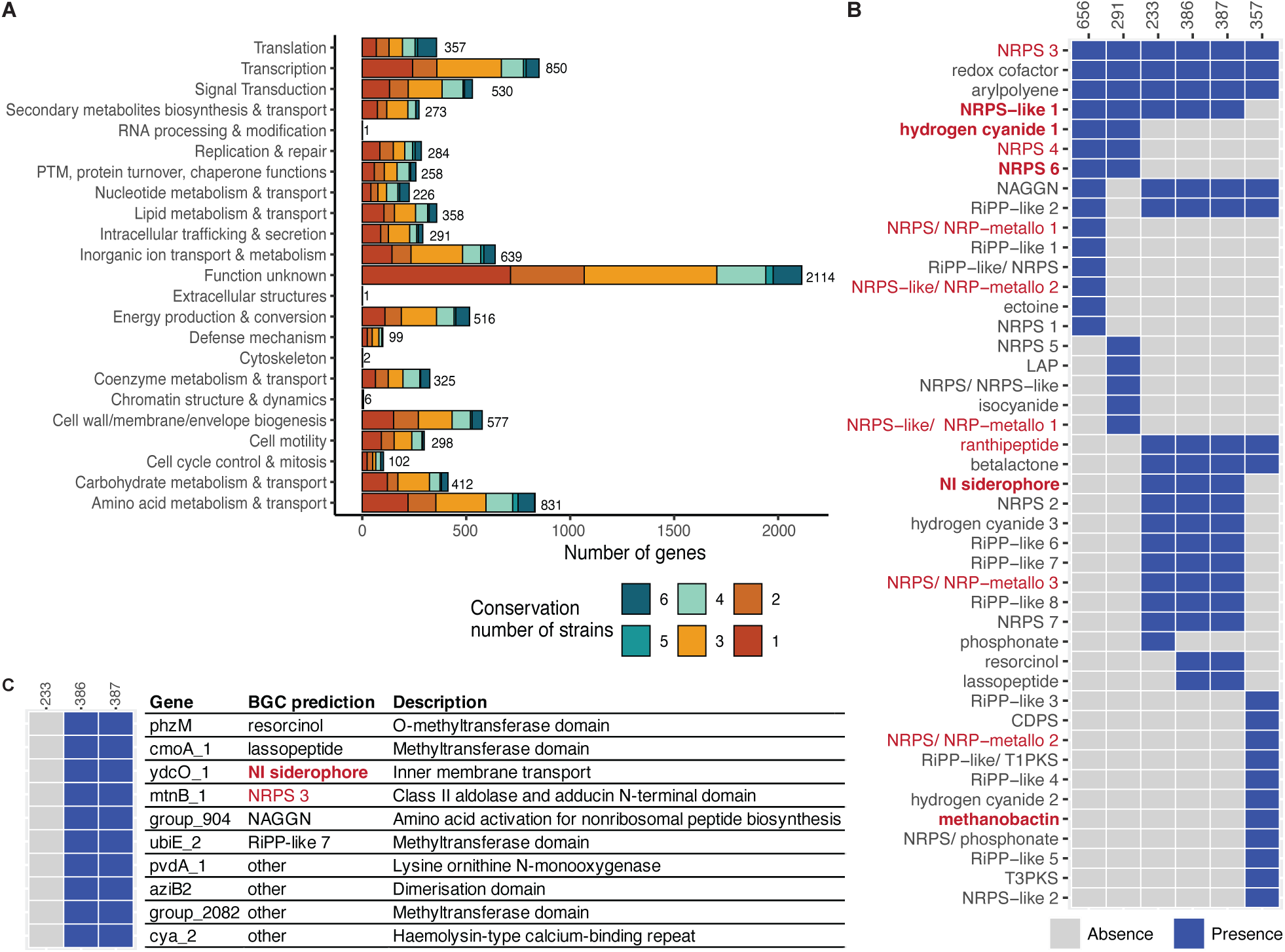
Genomic diversity and biosynthetic potential of six bioactive *Pseudomonas* isolates. (**A**) Functional classification of predicted proteins from these six *Pseudomonas* genomes, grouped by Cluster of Orthologous Gene (COG) categories and colored by the number of isolates in which they occur. (**B**) Presence-absence matrix of predicted BGCs across these six *Pseudomonas* genomes, with siderophore-encoding BGCs shown in red and previously unreported siderophore-encoding BGCs from *P. aeruginosa* shown in bold (metallo = metallophore). (**C**) Secondary metabolite-related genes unique to *P. aphyarum* isolates 386 and 387 as compared with isolate 233.

Secondary metabolism potential was further explored using antiSMASH (49) and BiG-SCAPE (50), revealing 44 predicted BGCs across the six sequenced genomes (Fig. 4B). The most frequently detected BGC classes included non-ribosomal peptide synthetases (NRPS), NRP-metallophores, and ribosomally synthesized and post-translationally modified peptides (RiPPs). Notably, three clusters were shared across all six *Pseudomonas* isolates, including NRPS, redox cofactor, and arylpolyene clusters, indicating their importance in this environment. Fourteen BGCs were predicted to encode siderophores, of which eight correspond to those known from *P. aeruginosa* (e.g., pyoverdine and enantio-pyochelin), which has been extensively studied for iron scavenging capacity. Six siderophore clusters were predicted to produce siderophores not yet reported from *P. aeruginosa*, representing potentially novel types with possible ecological advantages. Additionally, several BGC classes were unique to specific species, such as ectoine (*P. fontis*), linear azol(in)e-containing peptide (LAP) and isocyanide (*P. rubra*), and cyclodipeptide synthase (CDPS) and Type III polyketide synthase (T3PKS) (*P. idahonensis*), highlighting further functional diversity.

The genomes of three closely related *P. aphyarum* isolates (233, 386, and 387) showed over 99% average nucleotide identity (ANI) (Fig. S8) and nearly identical BGC profiles, while differing in bioactivity (Fig. 3). Notably, the BGCs encoding lassopeptide and resorcinol were absent in strain 233 but present in strains 386 and 387 (Fig. 4B), which both displayed biofilm disruption activity. Comparative genomic analysis identified ten secondary metabolite-related genes unique to strains 386 and 387 but absent in 233, several of which were annotated as methyltransferases and co-localized with distinct BGCs predicted using antiSMASH (Fig. 4C). Together, these differences suggest that small chemical modifications, such as methylation, could potentially modulate the differential bioactivity observed among these nearly clonal isolates.

### Highly abundant siderophores found across bioactive isolates

To identify small molecules produced by the six prioritized *Pseudomonas* strains that may confer antibacterial and biofilm inhibitory activities, chemical analyses of these isolates were performed using untargeted mass spectrometry to compare their metabolomic profiles. Annotation with the NPAtlas (45) and SIDERITE (46) databases resulted in 50 metabolomic features annotated as known siderophores, among which five features had distinctive MS2 profiles that were further validated as pyoverdine, tenacibactin B, pseudomonine, enantio-pyochelin, and hinduchelin A (Fig. 5A, S9). Pyoverdine and enantio-pyochelin have been previously reported from *P. aeruginosa* (Visca et al., 2007; Brandel et al., 2012), although neither has been reported from these new strains. Pyoverdine was genomically predicted by antiSMASH to be produced by all six strains, whereas enantio-pyochelin was genomically predicted for *P. idahonensis* 357 (Table S7). Metabolomics confirmed these genomic predictions, with pyoverdine shown to be produced by all six isolates, whereas enantio-pyochelin was only detected from strain 357. Pseudomonine (58) was predicted in the genome of strain 656 and was detected at high abundance from this strain. Tenacibactin B was also detected from strain 656 (59). Hinduchelin A was detected in strains 386, 387, and 357 at varying concentration (60). Of the evaluated six isolates, all those showing activity in the biofilm reduction assay [656, 386, 387, and 357 (Fig. 3B)] produced at least one unique siderophore that was not found in *P. aeruginosa*, namely tenacibactin B, pseudomonine, and hinduchelin A (Fig. 5A-B). Additionally, a higher abundance of hinduchelin A was associated with greater biofilm exculsion activity.

**Figure 5.**
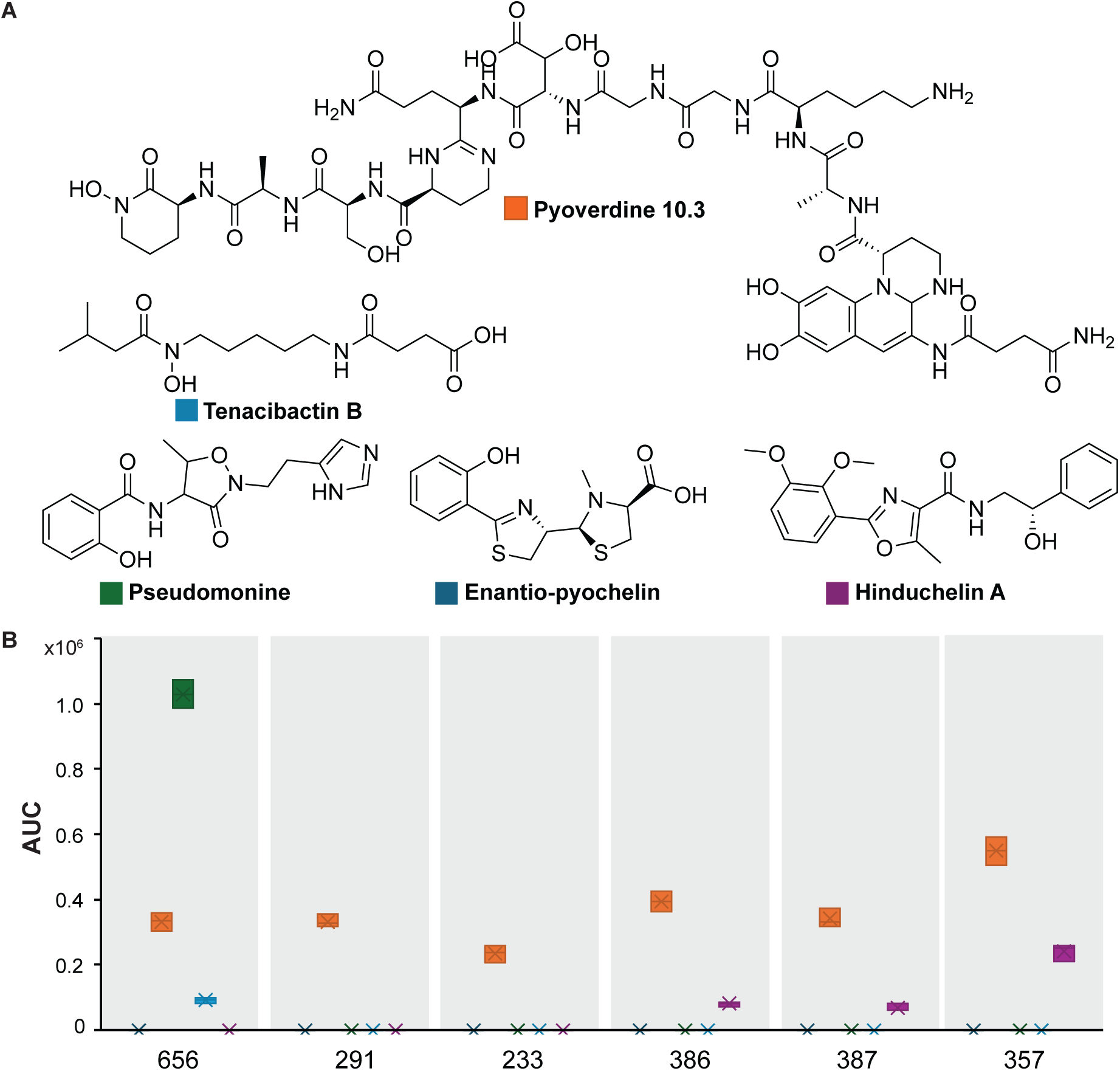
Siderophores were highly abundant in all prioritized aquaculture *Pseudomonas* strains. (**A**) Structures and (**B**) abundances [measured by area under the curve (AUC)] of five siderophores produced across all prioritized bioactive *Pseudomonas* strains. Pyoverdine was observed to be produced by all six strains, and hinduchelin A was produced by multiple strains, whereas pseudomonine, enantio-pyochelin, and tenacibactin B were each produced by a single strain.

### Co-culture with P. aeruginosa elicits production of secondary metabolites

Since extracts from aquaculture isolates 291 and 656 were more active against MSSA when co-cultured with *P. aeruginosa* (Fig. 3C), we conducted comparative metabolomics to putatively identify bioactive metabolites. To prioritize features of interest, metabolomics data was filtered for high-quality features using mpactR (44), with features from solvent and media blanks further filtered. Importantly, because these co-cultures included small quantities of *P. aeruginosa*, any features found in the *P. aeruginosa* monoculture were also filtered such that we prioritized features produced by the aquaculture strains.

Using volcano plots, 20 features were identified as significantly higher in the 291 + *P. aeruginosa* co-culture compared to 291 cultured alone (Fig. 6A, in red), eight of which were annotated via the NPAtlas database for bacterial compounds, with two of these having previously reported activity against MSSA (Fig. 6B). Viridenomycin, a 24-membered macrocyclic polyene lactam antibiotic first isolated from *Streptomyces viridochromogenes*, was previously reported to have antibiotic activity against *S. aureus* with an MIC of 0.5-1.0 μg/mL (61). Madurastatin C1, detected at a 2.8× higher abundance in the 291 + *P. aeruginosa* co-culture, is a pentapeptide siderophore first isolated from *Actinomadura* sp. and was reported to have antibiotic activity against *S. aureus* (62). Interestingly, multiple microcystins were produced in less abundance when 291 was co-cultured with *P. aeruginosa* (Fig. 6A, in blue).

**Figure 6.**
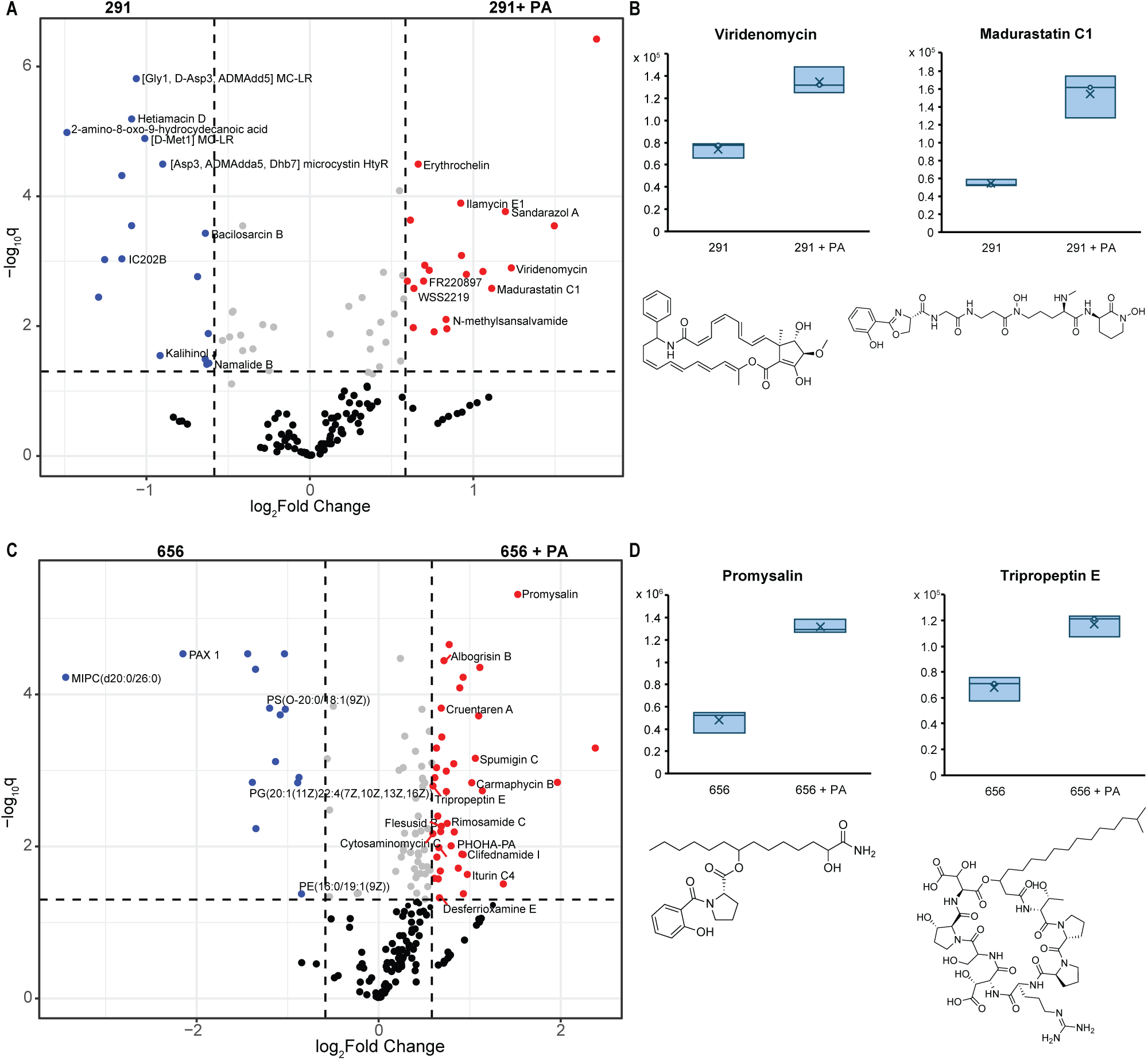
Multiple secondary metabolites with reported antibacterial and biofilm activity were produced in higher abundance when aquaculture strains were co-cultured with *P. aeruginosa*. Extracts from isolates 291 and 656 were both more active against MSSA when co-cultured with *P. aeruginosa* (Fig. 3C). Using comparative metabolomics, 20 metabolites were shown to be produced in higher abundance by isolate 291 when co-cultured with *P. aeruginosa* (PA, **A**). Several of these metabolites were annotated, including (**B**) viridenomycin, first isolated from *Streptomyces viridochromogenes* (61), and madurastatin C1, first isolated from *Actinomadura* sp. (62). When isolate 656 was co-cultured with *P. aeruginosa*, 37 metabolites were produced in higher abundance (**C**), including (**D**) promysalin, first isolated from *Pseudomonas putida* (63–65), and tripropeptin E, first isolated from *Lysobacter* sp. (66).

Using similar analyses, 39 features were significantly higher in isolate 656 + *P. aeruginosa* compared to isolate 656 alone, with 12 features annotated as bacterial metabolites using the NPAtlas database (Fig. 6C, in red). Among those annotated features, promysalin, a salicylate-containing antimicrobial first isolated from *Pseudomonas putida*, was produced by 656 at 2.8× higher abundance when co-cultured with *P. aeruginosa* (Fig. 6D). Promysalin was first reported to be selectively inhibitory against the *Pseudomonas* genus (63, 64), although it was later shown to exhibit broad-spectrum antibiotic activity, including against *S. aureus* (65).

Tripropeptin E, a non-ribosomal cyclic lipopeptide first isolated from *Lysobacter* sp. with reported activity against *S. aureus* (MIC of 0.78 μg/mL) (66), was produced at 1.6× higher abundance when 656 was co-cultured with *P. aeruginosa* (Fig. 6D). Interestingly, four lipids from different classes were detected at lower abundances when 656 was co-cultured with *P. aeruginosa* (Fig. 6C, in blue), suggesting there may be some disruption to lipid biosynthesis and/or metabolism that may confer differential activity.

To determine if there was an overall effect on the aquaculture isolates when co-cultured with *P. aeruginosa*, a molecular network was constructed to investigate related groups of features that increased in abundance across multiple isolates when co-cultured with *P. aeruginosa* (Fig. S10). In this network, each node represents an mpactR filtered metabolomic feature. A bioactivity ranking score was calculated for each feature, with higher scores indicating greater abundance in co-cultures that exhibited stronger activity (Fig. 3C). Larger nodes represent metabolomic features produced by multiple isolates in higher abundance under co-culture that may be indicative of relevant biological function. For example, a triple-charged, peptide-like feature with the mass of 1946.9828 was produced in higher abundance in all four isolates (strains 656, 291, 386, and 387) that showed increased activity against MSSA when co-cultured with *P. aeruginosa* (Fig. S10C). Thus, there are opportunities for discovery of new bioactive molecules from these strains for future studies.

## Discussion

Our goal was to identify pathogen-inhibiting bacterial isolates that are part of the natural biofilm community within a rainbow trout aquaculture facility with the long-term objective of developing novel approaches to reduce the presence of pathogens in aquaculture settings. This initial work produced promising candidates that inhibit the growth of important aquaculture pathogens. From our initial screen of 510 isolates, we identified 29 isolates that inhibited aquaculture pathogens. This investigation led to the identification of a group of recently described *Pseudomonas* species (56) that were shown to inhibit a suite of fish and human pathogens *in vitro* through agar plate-based screens and exclude a fish pathogen from establishing itself in a biofilm.

We were able to identify 510 isolates from aquatic biofilms to the species level by leveraging full-length 16S rRNA gene sequencing by long-read sequencing using PacBio. Using barcodes and multiple SMRTbells, we were able to identify these strains on a single PacBio sequencing run and significantly reduce potential costs for identification. When comparing the abundance and presence of specific phyla from our isolate catalogue to 16S metagenomic survey data generated from this same location (15), it is evident that specific groups were missed.

Proteobacteria and Bacteroidota are the predominant phyla in these biofilm communities (comprising >75% of the total composition) and represented the majority of groups we isolated. However, some of the less abundant phyla, such as Verrucomicrobiota (∼10% of community), Myxococcota (∼5%), Bdellovibrionota (∼5%), and Planctomycetota (∼2%), were not recovered in our culturing effort. These groups likely have important ecological roles in these biofilm communities, and their isolation would prove valuable for investigating these roles. Other groups, such as OM190 within the Planctomycetota, were completely uncultured but comprise a significant portion of this community. Alternate specialized media types and growth conditions, including liquid media, anaerobic culturing, co-culturing, and media recipe construction based on the nutrient profile of the spring water used at the farm could be used to attempt to culture some of these lesser studied, but very interesting groups.

The inhibitory screens identified a group of 29 isolates with inhibitory ability against one or more fish or human pathogen strains. These isolates were almost entirely *Pseudomonas*, which is not particularly surprising since *Pseudomonas* has been found to produce a wide variety of secondary metabolites with an assortment of functions (67). However, the source location being aquatic biofilms, where competition and evolution of advantageous traits is arguably at its apex (67–69), makes these isolates interesting despite being from an otherwise well-studied genus. *P. rubra* and *P. aphyarum* isolates performed the best across the fish pathogen inhibition tests (inhibiting 24 or more of the 27 tested strains), with isolates 386 and 387 inhibiting every tested fish pathogen. These strains were also able to inhibit some of the ESKAPE pathogens, with a subset additionally being effective at excluding *F. columnare* from a pre-formed biofilm. For the biofilm exclusion assays against *F. columnare*, *P. aphyarum* 386 and 387 were some of the best performing strains in addition to *P. idahonensis* 357.

The comparative genomic analysis of closely related strains revealed interesting differences in the potential of these strains to produce bioactive molecules. The intra-species variation revealed that *P. idahonensis* 357 contains one additional bacteriocin cluster and one additional CDPS cluster while *P. idahonensis* 1048 contains two additional NRPS-associated clusters. In the inhibition data, isolate 357 inhibited a much wider diversity of *A. salmonicida* and *Y. ruckeri* strains along with more ESKAPE pathogens than isolate 1048, possibly due to these additional bacteriocin and CDPS clusters. *P. fontis* 656 inhibited very few *Y. ruckeri* and *F. columnare* strains while *P. fontis* 681 inhibited almost all of them. Isolate 681 has three additional NRPS clusters that may contribute to the observed differences between isolates within the same species. *P. aphyarum* 386 and 387 have two unique metabolite clusters (lassopeptide and resorcinol) that *P. aphyarum* 233 lacks. In the biofilm exclusion assay, 233 performed far worse than 386 or 387 despite plate inhibition results being relatively similar. It is possible that one or both secondary metabolite clusters somehow improve biofilm formation by the *Pseudomonas* isolates, actively functioning to exclude *F. columnare* from the biofilm, or are only expressed when these isolates are existing in this lower oxygen biofilm setting (as opposed to on a plate). Lassopeptides are RiPPs that have been noted as having variable antimicrobial effects (71), with only one report of a *Pseudomonas*-associated lassopeptide from a deep-sea organism (72). Resorcinol is used as a topical antiseptic that has been shown to be produced by a *Pseudomonas chlororaphis* strain for agricultural biocontrol of a fungal pathogen (73). Deeper genomic comparison at the gene level revealed that 10 orthologous genes involved in secondary metabolite biosynthesis and transport were present in 386 and 387 while absent in 233, four of which were putative methyltransferases. Differences in the ability of strains to inhibit the growth of pathogens were mirrored by differences in genomic content.

Chemical analyses of the prioritized subset of six aquaculture isolates, including some active in both antimicrobial and biofilm assays as well as and some with differential activity, identified highly abundant siderophore production. Siderophores are high-affinity iron-chelating compounds produced by microorganisms to scavenge iron from the environment, a critical nutrient often limited in availability. These molecules have been extensively studied for their role in microbial virulence, competition, and biofilm dynamics (74, 75). In our study, siderophore production was confirmed using untargeted metabolomics in several bioactive *Pseudomonas* strains. This result supports a link between siderophore production and biofilm exclusion capabilities in *Pseudomonas*, which suggests that interfering with iron acquisition may be a robust strategy to combat pathogenic biofilm formation and mediate community interactions.

Multiple siderophores were detected across the bioactive *Pseudomonas* isolates, underscoring their potential role in biofilm disruption. Pyoverdine, a widely studied siderophore produced by *P. aeruginosa*, was found in all six strains at varying concentrations, emphasizing its central role in iron competition and biofilm stability (76). Hinduchelin A, previously reported in *Streptoalloteichus hindustanus* (60), was detected in our three most biofilm-active isolates, raising the possibility of horizontal gene transfer into *Pseudomonas* and illustrating potential evolutionary plasticity of siderophore biosynthetic pathways. Interestingly, the remaining annotated siderophores (enantio-pyochelin, pseudomonine, and tenacibactin B) demonstrated strain-specific production, suggesting specialized ecological functions and adaptive value for niche colonization or competition under variable environment conditions, such as varying pH or iron availability (77). For example, enantio-pyochelin was unique to *P. idahonensis* isolate 357, while pseudomonine and tenacibactin B were exclusive to *P. fontis* isolate 656. These findings align with established knowledge regarding the biofilm-disrupting capabilities of siderophores but also identify new siderophore candidates that may hold unique biotechnological application.

The rich diversity of siderophore profiles from our biofilm isolates suggests a balance between widespread and specialized iron-acquisition strategies. This diversity not only shapes microbial competition, but may also reflect the influence of siderophore “cheating,” a phenomenon where microorganisms utilize siderophores produced by other members of the community to fulfill their own iron requirements, a critical aspect of microbial interactions within biofilm communities (78). This strategy allows non-producers to access iron without the metabolic cost of siderophore biosynthesis (78–80), generating evolutionary pressure for the emergence of novel or less exploitable siderophores (81, 82). In the context of our study, strains possessed a combination of shared and strain-specific siderophores, suggesting a complex dynamic interplay between cooperative and competitive strategies within the biofilm environment. Detailed knowledge of the affinity of the different siderophores and the prevalence of their receptors would aid in an understanding of the need to express multiple siderophores.

Understanding the contribution of siderophore cheating to biofilm ecology will be important for developing strategies that leverage probiotic isolates to effectively disrupt harmful biofilms in aquaculture and beyond.

Co-culture experiments of our aquaculture isolates with *P. aeruginosa* revealed an inducible response for secondary metabolite production, particularly for antibacterial compounds, when challenged with a competing microorganism, consistent with the concept of microbial interactions driving bioactive compound production as a defensive strategy (83). The observed increase in antibiotic activity of co-culture extracts against MSSA illustrates the dynamic metabolic adaptation driven by competitive pressure and highlights the potential of co-culture methodologies for discovering novel bioactive compounds. Several key metabolites, including viridenomycin, madurastatin C, promysalin and tripropeptin E, were putatively identified at higher levels in co-cultures, suggesting their involvement in competitive inhibition and inhibitory mechanisms against *P. aeruginosa*. These findings underscore the value of inhibition assays involving live microbial interactions, which may reveal bioactive compounds that could be overlooked using conventional extract-only screens.

Given the demonstrated bioactivity of these *Pseudomonas* strains against important pathogens, they hold promise as sources for novel antimicrobial discovery. While further testing into feasibility, efficacy, and safety is needed, an interesting strategy may be to seed trout raceway biofilms with these natively-sourced bacteria to exclude pathogens within these biofilms. This approach may provide a practical solution to mitigating disease burdens and reducing reliance on conventional antibiotics in aquaculture farming.

## Data Availability

Genome assemblies have been deposited in NCBI under BioProject PRJNA835116. Metabolomic data sets used in this work were deposited in the Mass Spectrometry Interactive Virtual Environment repository MassIVE (https://massive.ucsd.edu/ProteoSAFe/static/massive.jsp) under accession number MSV000098842.

## Author Contributions

J.N.T.N and T.T. contributed equally to this work. J.N.T.N, T.T., M.J.B., and J.G. designed the research. J.N.T.N, T.T., K.R.M., H.D., and J.M. performed experiments and analyzed data. J.N.T.N, T.T., M.J.B., and J.G. wrote and edited the paper.

## Acknowledgements

This work was supported by the U.S. Department of Agriculture [8082-32000-007-00-D].

We thank the University of Delaware DNA Sequencing and Genotyping Center for performing the PacBio sequencing for the initial 16S rRNA gene identifications. We thank the University of Connecticut Microbial Analysis, Resources, and Services (MARS) facility for performing the Illumina sequencing for the genomes. We thank Drs. Timothy Welch, and Hugh Cai for supplying some of the pathogen strains. We thank Dr. Mark Driscoll, Dawn Gratalo, and Eric Jackson from Intus Biosciences (formerly Shoreline Biome) for support with 16S rRNA gene library preparations and data processing. We thank Hannah Boesger for initial cultures and extraction of prioritized isolates.

